# Toward asleep DBS: Cortico basal-ganglia neural activity during interleaved propofol/ketamine sedation mimics NREM/REM sleep activity

**DOI:** 10.1101/2020.12.14.422637

**Authors:** Jing Guang, Halen Baker, Orilia Ben-Yishay Nizri, Shimon Firman, Uri Werner-Reiss, Vadim Kapuller, Zvi Israel, Hagai Bergman

**Author notes:** corresponding author, Edmond and Lily Safra Center for Brain Sciences, The Suzanne and Charles Goodman Brain Sciences Building, 3. Equal contribution (Jing Guang, Halen Baker, Orilia Ben-Yishay Nizri, Shimon Firman).

## Abstract

Deep brain stimulation (DBS) is currently a standard long-term treatment for advanced motor symptoms in Parkinson’s disease (PD). In an effort to enable DBS under sedation, asleep DBS, we characterized the cortico-basal ganglia neuronal network of two non-human primates under propofol, ketamine and interleaved propofol-ketamine (IPK) sedation. Further, we compared these sedation states in the healthy and Parkinsonian condition to those of healthy sleep. Ketamine increases high frequency power and synchronization while propofol increases low frequency power and synchronization in polysomnography and neuronal activity recordings. Thus, ketamine does not mask the low frequency oscillations used for physiological navigation toward basal ganglia DBS targets. The brain state under ketamine and propofol mimicked rapid eye movement (REM) and Non-REM (NREM) sleep activity, respectively, and the IPK protocol imitates the NREM-REM sleep cycle. These promising results are the first step towards asleep DBS with non-distorted physiological navigation.

---

It is estimated that 10 million people worldwide suffer from Parkinson’s disease (PD). While dopamine replacement therapy offers a good short-term solution, after five to 10 years severe side effects emerge. Currently, one of the most promising long-term treatments available is Deep Brain Stimulation (DBS). The DBS procedure aims to surgically implant leads that enable high-frequency stimulation of specific nuclei in the basal ganglia. The most common targets are the subthalamic nucleus (STN) and the internal segments of the globus pallidus (GPi). Behaviorally, this results in a vast improvement in PD symptoms^1–4^. DBS is not only used for PD patients but additionally for patients with dystonia, essential tremor and psychiatric disorders^5–8^ suggesting that enhancement of this technique would be quite far reaching. To accurately reach the target of the DBS lead, a neural navigation system that requires brain electrophysiological signals from the awake patient may be used^9,10^. Many patients avoid DBS therapy due to the fear of undergoing awake brain surgery, leaving a wide gap for therapeutic improvement.

Propofol is currently the most commonly used sedative hypnotic drug in clinical anesthesia. In parkinsonian patients, propofol might cause dyskinesia (probably due to paradoxical excitation) or abolish tremor^11^. Nevertheless, many publications support the use of propofol in PD patients because of its fast onset and short duration of action. On the other hand, studies in human subjects undergoing DBS procedure demonstrated a significant modulation of the neuronal discharge of basal ganglia DBS targets in response to propofol sedation^12,13^. Changes in cortical activity have also been reported under propofol^14^. These changes in cortex and STN/GPi discharge rate and pattern might interfere with the detection of the DBS targets. However, the short washout period of propofol may offer an ideal solution when used interleaved with a non-disruptive sedative.

Ketamine, a dissociative agent, is less commonly used in neuro-anesthesia due to its reputation as causing increases in cerebral blood flow and metabolism, increased cranial pressure (ICP) and frightening hallucinations. However, recent studies revealed that ketamine does not affect cerebral metabolism^15–17^ or ICP^18^, and that sub-anesthetic doses of ketamine are associated with good subjective experience^19^. Ketamine has analgesic effects and has in fact shown promise as a treatment for depression^20^. In animal models, ketamine increases the spontaneous gamma and ultra-slow oscillations in cortical and basal ganglia nuclei^21–26^.

Akinesia of Parkinson’s patients and animal models is associated with beta oscillations^27^, while Parkinson’s therapies (L-DOPA and DBS) are associated with gamma oscillations^28,29^. This leads us to believe that ketamine sedation may be less disruptive to the navigation system that uses beta oscillations for detection of DBS targets and subdomains and potentially a good choice for use in DBS surgeries.

The relationship between sleep and sedation is multifaceted and is still not completely understood. Due to variations in the mechanisms of action of sedative agents, there is a wide range of reported similarities and differences between natural sleep and any given sedative medication^30,31^. For patients already ill, good quality sedation is essential to their wellbeing. Thus, mimicking natural sleep with sedation could greatly benefit them both during and following surgeries. To this end, we aimed to test the effects of propofol and ketamine sedation and an interleaved propofol-ketamine (IPK) sedation protocol on the neural activity in the cortex-basal ganglia circuit. We hypothesize that propofol would enable the more invasive stages of DBS procedures (like drilling of the burr-hole) and due to its fast action its effects on the basal ganglia would not be persistent. In a complementary way, ketamine may be undisruptive to the neural navigation system while still providing the necessary dissociative and analgesic effects needed during this less invasive stage of the surgery. If correct, this will allow sedation to be safely given during navigation and lead implantation in DBS surgery and greatly improve the experience of these patients.

## Results

The experiments were performed on two female African green monkeys (*Chlorocebus aethiops sabaeus* (vervet), weight: ∼4 kg). All experimental protocols were conducted in accordance with the National Institutes of Health *Guide for the Care and Use of Laboratory Animals* and The Hebrew University of Jerusalem guidelines for the use and care of laboratory animals in research. Prior to the surgery, the non-human primates (NHPs) were introduced to the recording room and trained to sit and sleep in a specially made primate chair. The first surgery consisted of a craniotomy and implantation of a recording chamber, head holder and EEG electrodes. The second surgery involved the implantation of a subcutaneous ported central venous catheter. The experiments included neural recording (Figure 1a) while the animal was under sedative medications and naturally sleeping. The recordings were carried out before and after systemic treatment with 1-methyl-4-phenyl-1,2,3,6-tetrahydropyridine (MPTP) neurotoxin and induction of Parkinson’s symptoms (Figure 1b and Table S1) and with careful monitoring of the vital signs (Figure 1c). After the end of the recording the NHP’s were rehabilitated, and moved to the Israel Primate Sanctuary (www.ipsf.org.il).

**Fig 1:**
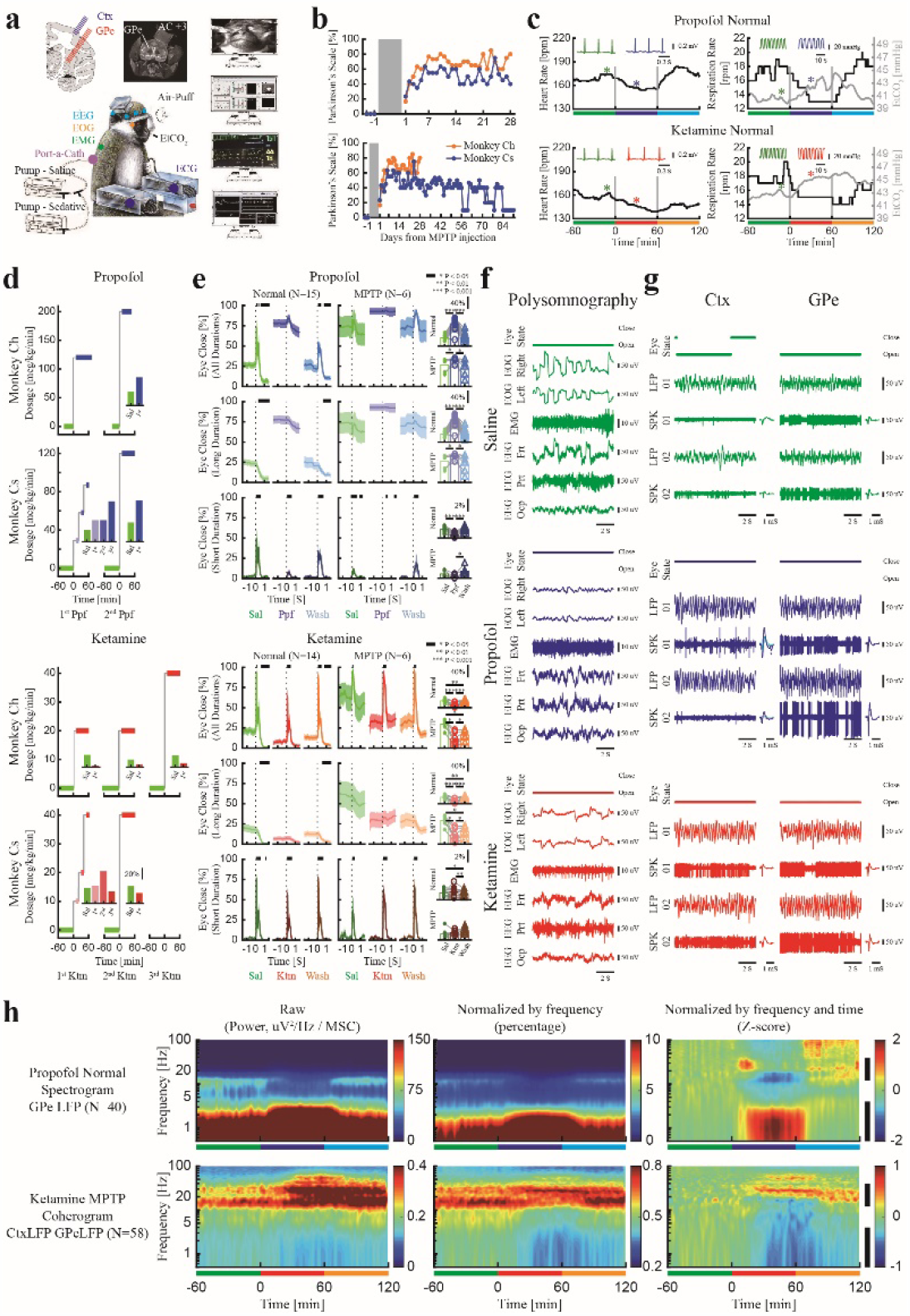
Experimental procedures. **a**. Recording setup. **b**. Parkinsonian symptoms after MPTP-treatment. Upper: recording period, lower: recording and recovery periods. **c**. The heart rate, respiration rate and end tidal CO_2_ from example sessions of propofol (upper) and ketamine (lower) procedures. The inset plots show examples of ECG and end tidal CO_2_ (from time locations marked with asterisks). Green: saline baseline, blue: propofol sedation, red: ketamine sedation and cyan: propofol saline washout, orange: ketamine saline washout. **d**. Titration process. Inset shows average proportion of time with eyes closed. **e**. Eye closure proportion for each stage before (normal, left) and after MPTP treatment (right). From upper to lower, each row shows proportion of eyes closed for all durations, long and short duration (blink, < 1 S). Shaded area shows SEM. Top black bar represents significant difference in eye closure compared with period before time 0 (p<0.05, Wilcoxon rank-sum test). The scatter and bar plots show average (over time) eye closure proportions. P-value is given in Table S2, Wilcoxon signed-rank test. **f**,**g**. Examples of 10 second traces of polysomnography, LFP/SPK of Ctx/GPe during saline baseline (upper, green), propofol (center, blue) and ketamine (lower, red) and average spike waveform. **h**. Normalization process of spectrogram and coherogram. Vertical black bar shows the frequency range for the high/low power and synchronization differences. Ctx, cortex. ECG, electrocardiogram. EEG, electroencephalogram. EMG, electromyography. EOG, electrooculography. EtCO_2_, end tidal CO_2_. Frt, frontal. GPe, globus pallidus external segment. Ktm, ketamine. LFP, local field potential. MSC, magnitude-squared coherence. NHPs, non-human primates. Ocp, occipital. Ppf, propofol. Prt, parietal. Sal, saline. SPK, spiking.

First, titration sessions were performed for each sedative drug and each NHP to establish moderate sedation dose (Figure 1d). Sedation sessions consisted of one hour of saline baseline followed by one hour of sedation (propofol/ketamine) and finally one hour of saline washout. Eye open/close state and blinking response to air puff stimulation to the eye (Figure 1e, Table S2), polysomnography, including electroencephalogram, electrooculogram, and electromyogram (EEG, EOG, EMG, Figure 1f) and local field potential and spiking activity (LFP and SPK, Figure 1g) from the frontal cortex and globus pallidus external segment (GPe) were recorded during all three phases of each session.

Currently, DBS physiological navigation systems use the independent parameters of discharge rate (or total power of spiking activity) and discharge pattern (e.g. power spectrum densities normalized to the total power). The total power of the spiking activity is often used for detection of the borders of the structures, while the spectral signature enables the discrimination of the target subdomains (STN and GPi motor domains are characterized by theta and beta oscillations). Both parameters can be affected by sedation agents. Firing rate and total power of spiking activity were modulated during propofol and ketamine sedation (Figure S1). Here, we will mainly discuss the effect of these sedation agents on the discharge pattern. Power spectrum densities and pairwise coherence were calculated as a function of time (spectrograms and coherograms) for all EEG, LFP and SPK recordings and normalized by frequency (i.e. by their total power/coherence) and by the saline baseline period (by frequency and by time, Figure 1h).

### Propofol and Ketamine differentially modulate power and synchronization

To establish the individual characteristics of propofol and ketamine in the healthy and parkinsonian conditions, we investigated power spectrum changes during the sedation period for each recording target and modality (Figure 2, Figure S2). This revealed that in comparison to the saline baseline in all recording modalities (EEG, EMG, EOG, LFP and SPK) and targets (cortex and basal ganglia) propofol sedation increased low frequency power. Particularly, propofol sedation boosted delta frequency power, in EEG and LFP of both the frontal cortex and GPe (Figure 2a,b). On the other hand, ketamine sedation decreased low frequency power and increased high frequency power, particularly gamma frequency power, in all modalities (Figure 2c,d). MPTP treatment and the emergence of Parkinson’s symptoms resulted in changes in the neural activity (Figure S3). Nevertheless, there were no significant differences between the sedation effects in the parkinsonian condition and the healthy condition (Figure 2a-d).

**Fig 2:**
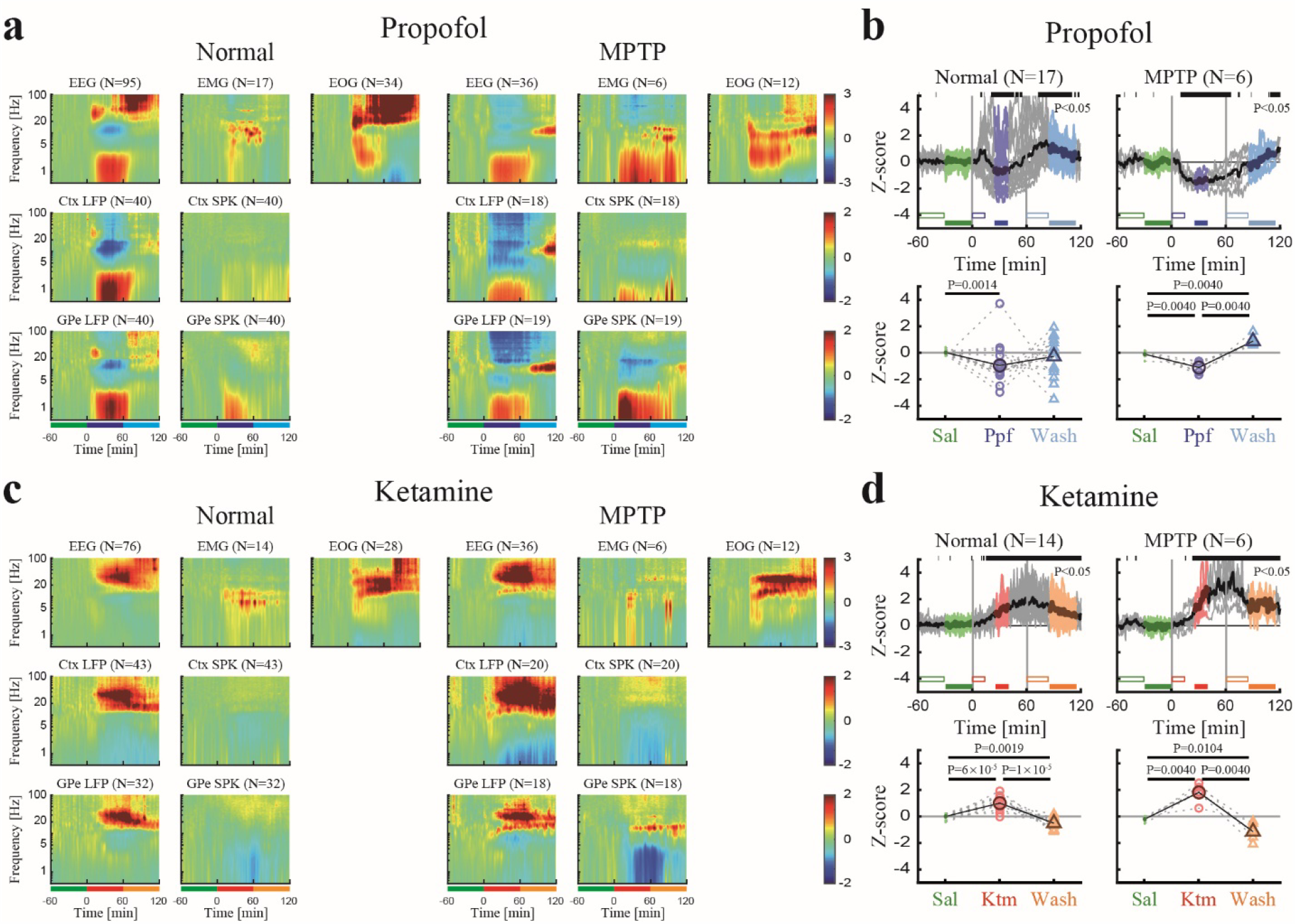
Propofol and ketamine increase low and high frequency power, respectively, in polysomnography and cortico-basal ganglia neural activity. **a**. The normalized power spectrograms of EEG/EMG/EOG (1 ^st^ row), Ctx LFP/SPK (2 ^nd^ row) and GPe LFP/SPK (3 ^rd^ row) during saline baseline, propofol sedation, saline washout before (left) and after (right) MPTP-treatment. Lower bar represents 1-hour time periods of saline baseline (green), propofol sedation (blue) and saline washout (cyan). **b**. Upper. The normalized high/low power difference (averaged through all spectrograms within one session) between spectral power at high frequency (12-40 Hz) and low frequency (0.5-4 Hz) domains before (left) and after MPTP-treatment (right). Top black bar represents significant difference compared to saline (p<0.05, Kruskal–Wallis test). Lower. The change from baseline of the normalized high/low power difference during saline baseline (upper, green minus empty green), propofol sedation (upper, blue minus empty blue), saline washout (upper, cyan minus empty cyan) before (left) and after (right) MPTP-treatment. P-value is given, Kruskal–Wallis test. **c, d**. Same conventions as **a, b**. during ketamine. Color represents saline baseline (green), ketamine sedation (red) and saline washout (orange). Number of sessions (b, d) and sites (a, c) is given for both monkeys in each subplot. Abbreviations as in Fig.1.

Similarly, we performed an exploration of the pair-wise synchronization (coherence) of EEG, cortex and basal ganglia LFP and spiking activity. We found propofol sedation increased low frequency synchronization while ketamine increased high frequency synchronization in, and between the frontal cortex and basal ganglia in both the healthy and parkinsonian conditions (Figure 3, Figure S4).

**Fig 3:**
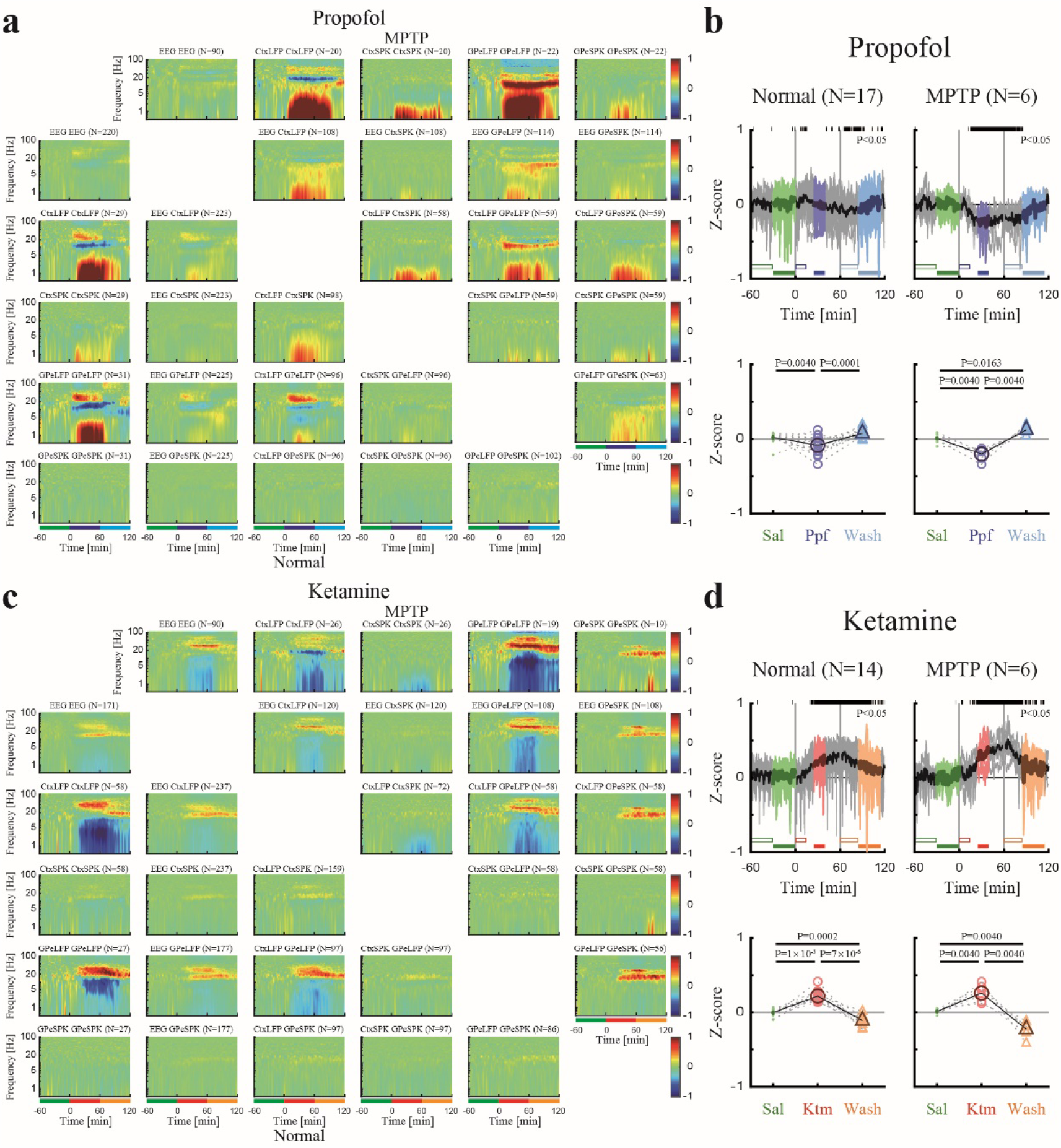
Propofol and ketamine increase and decrease low frequency synchronization, respectively, in cortico-basal ganglia neuronal activity. **a**. The normalized magnitude-squared coherograms of all pairs of EEG, Ctx LFP/SPK and GPe LFP/SPK during saline baseline, propofol sedation, saline washout before (lower left) and after MPTP-treatment (upper right). The 1^st^ row/column shows coherograms within the same signal type (e.g., EEG to EEG), other subplots show coherograms between different signal types. Lower bar represents 1-hour time periods of saline baseline (green), propofol sedation (blue) and saline washout (cyan). **b**. Upper. The normalized high/low synchronization difference (averaged through all coherograms within one session) between high frequency (12-40 Hz) and low frequency (0.5-4 Hz) coherence domains before (left) and after (right) MPTP-treatment. Top black bar represents significant difference compared to saline (p<0.05, Kruskal–Wallis test). Lower. The change from saline baseline of the normalized high/low synchronization difference during saline (upper, green minus empty green), propofol sedation (upper, blue minus empty blue), saline washout (upper, cyan minus empty cyan) before (left) and after MPTP-treatment (right). P-value is given, Kruskal–Wallis test. **c, d**. Same conventions as **a, b**. during ketamine. Color represents saline baseline (green), ketamine sedation (red) and saline washout (orange). Number of pairs (a, c) and sessions (b, d) is given for both monkeys for each subplot. Abbreviations as in Fig.1.

### Propofol mimics NREM sleep and ketamine mimics REM sleep

In order to examine the common properties of sleep stages and the two sedative agents, we calculated the spectrograms and coherograms of different sleep stages (Figure S5) for each modality. This revealed that in EEG, LFP and SPK cortical and basal ganglia recordings NREM sleep increased low frequency power while REM sleep increased high frequency power (Figure 4a,b, Figure S6a). Not only were the power analyses of sedation and sleep effects similar but the synchronization analyses (Figure 4c,d, Figure S6b) also showed common characteristics. NREM showed an increase in low frequency synchronization of the neural activity while REM showed a decrease in low frequency synchronization and an increase in high frequency synchronization. However, one way in which propofol and ketamine do not mimic the NREM/REM cycle is in the EMG and EOG recordings (Figure S7). Ketamine does not seem to induce atonia (full muscle relaxation) as seen in REM sleep nor does it cause the namesake rapid eye movements of REM sleep (Figure 1f vs. Figure S5c).

**Fig 4:**
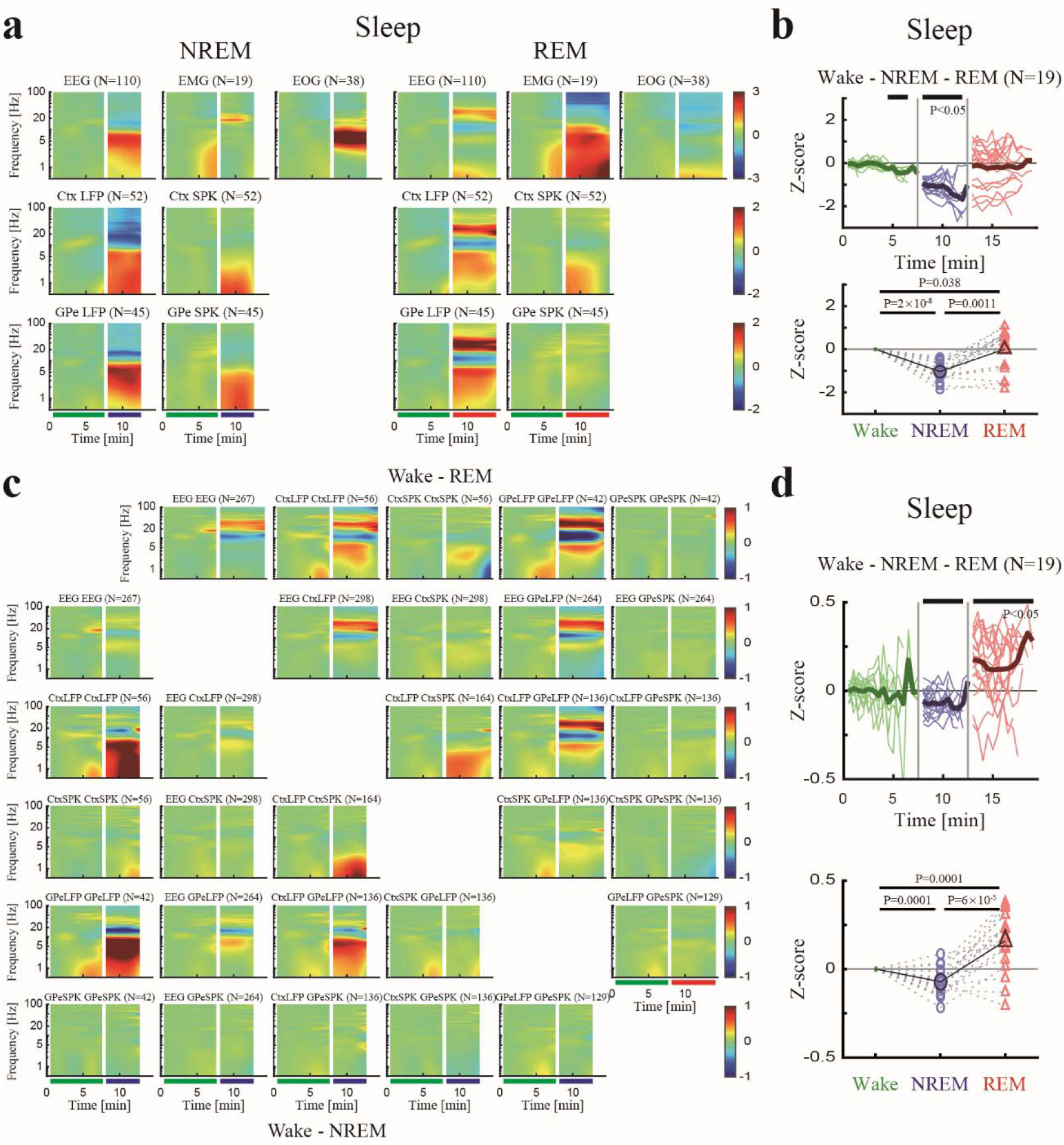
Polysomnography and neural activity during natural NREM and REM sleep show increased low frequency power/synchronization and increased high frequency power/synchronization, respectively. **a**. The normalized power spectrograms of EEG/EMG/EOG (1^st^ row), Ctx LFP/SPK (2 ^nd^ row) and GPe LFP/SPK (3 ^rd^ row) during NREM (left) and REM (right). Lower bar represents time periods of wake (green), NREM (blue) and REM (red). Data was averaged over different segments and different time periods are not necessarily sequential. **b**. Upper. The normalized high/low power difference (averaged through all spectrograms within one night) between spectral power at high frequency (12-40 Hz) and low frequency (0.5-4 Hz) during wake, NREM and REM. Top black bar represents significant difference compared to wake (p<0.05, Kruskal–Wallis test). Lower. The averaged normalized high/low power difference during wake, NREM and REM. P-value is given, Kruskal–Wallis test. **c**. The normalized magnitude-squared coherograms of all pairs of EEG, Ctx LFP/SPK and GPe LFP/SPK during NREM (lower left) and REM (upper right). The 1^st^ row/column shows coherograms within same signal type (e.g., EEG to EEG), other subplots show coherograms between different signal types. Lower bar represents time periods of wake, NREM and REM. **d**. Same conventions as **b** for synchronization difference. Number of sites (a), pairs (c) and nights (b, d) is given for both monkeys for each subplot. Abbreviations as in Fig.1. NREM, non-rapid eye movement sleep. REM, rapid eye movement sleep.

### Interleaved propofol-ketamine imitates the sleep cycle

Finally, to test if the effects of propofol and ketamine remain when simulating a natural sleep cycle we created an interleaved propofol and ketamine sedation session in which propofol and ketamine were administered such that a propofol-ketamine-propofol-ketamine-propofol cycle was established. In this regimen, propofol was administered for 40 minutes, directly followed by ketamine sedation for 20 minutes.

Similar to the changes observed in natural sleep from NREM to REM, the interleaved propofol-ketamine (IPK) sedation protocol followed the changes in spectrum and synchronization shown in the neural activity during propofol and ketamine sedations alone. Low frequency power and synchronization increases during propofol followed by high frequency power and synchronization increases during ketamine (mimicking NREM and REM respectively, Figure 5, Figure S8).

**Fig 5:**
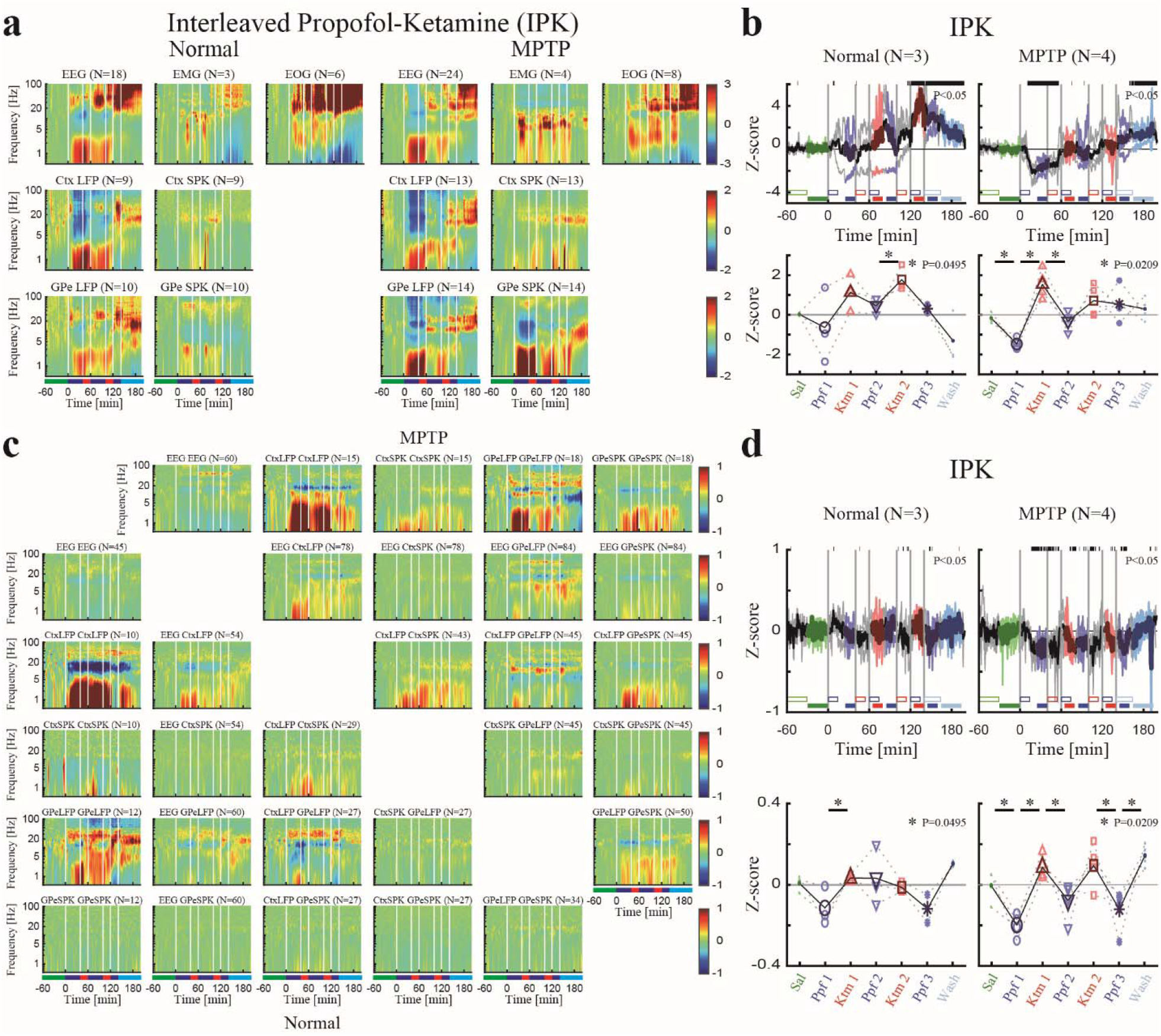
Interleaved propofol-ketamine (IPK) sedation protocol shows similar trend of power/synchronization changes as sedation induced by propofol and ketamine alone. **a**. The normalized power spectrograms of EEG/EMG/EOG (1^st^ row), Ctx LFP/SPK (2 ^nd^ row) and GPe LFP/SPK (3 ^rd^ row) during IPK before (left) and after (right) MPTP-treatment. Lower bar represents time periods of saline baseline (green), propofol sedation (blue), ketamine sedation (red) and saline washout (cyan). **b**. Upper. The normalized high/l ow power difference (averaged through all spectrograms within one session) between high frequency (12-40 Hz) and low frequency (0.5-4 Hz) mean power before (left) and after (right) MPTP-treatment. Top black bar represents significant difference compared to saline (p<0.05, Kruskal–Wallis test). Lower. The change from baseline of the normalized high/low power difference during saline (upper, green minus empty green), propofol sedation (upper, blue minus empty blue), ketamine sedation (upper, red minus empty red), saline washout (upper, cyan minus empty cyan) before (left) and after (right) MPTP-treatment. P-value is given, Kruskal–Wallis test. Statistics applied to adjacent sedation stages. **c**. The normalized coherograms of all pairs of EEG, Ctx LFP/SPK and GPe LFP/SPK during IPK before (lower left) and after (upper right) MPTP-treatment. The 1^st^ row/column shows coherograms within same signal type (e.g., EEG to EEG), other subplots show coherograms between different signal types. **d**. Same conventions as **b** for synchronization difference. Number of sites (a), pairs (c) and sessions (b, d) is given for both monkeys for each subplot. Abbreviations as in Fig.1.

## Discussion

This study aimed to characterize the electrophysiological changes during moderate ketamine sedation, propofol sedation, interleaved propofol-ketamine (IPK) sedation protocol and natural sleep using polysomnography and neuronal activity of the cortico-basal ganglia neuronal network. In EEG and LFP recordings from the frontal cortex and the basal ganglia, ketamine was found to increase high frequency power and synchronization while propofol was found to increase low frequency power and synchronization. These results were corroborated in the IPK protocol. The dynamic between propofol and ketamine was found to be a fast one and to mimic the relationship of NREM and REM natural sleep.

Classically, the target for DBS for PD is the STN^1^ while for dystonia it is the GPi^5^. Here, due to recording and time constraints we felt it was most appropriate to record and characterize the GPe. The three-layer model of the basal ganglia^32,33^ positions the GPe in the middle layer, receiving projections from the STN and projecting back to it as well as projecting to the GPi.

GPe recoding therefore permits us to best characterize the basal ganglia activity as a whole and to generalize our findings to both the STN and GPi. Clinically sedation level is often assessed using EEG characteristics and subjective measures such as response to verbal command or physical touch. Here, we used eye open/close states and blinking response to air puff to the eye, analogous to physical touch, to ascertain moderate sedation. Still, this could potentially be misleading as response may depend not only on sedation level but also mood or sleepiness. Nevertheless, we suggest that with constant monitoring and adjustment in the clinical setup, moderate sedation by propofol and ketamine is achieved without a problem.

At present, there is tension between patient comfort and precise implantation of the lead during DBS surgery. Only a small fraction of eligible PD patients choose to undergo DBS surgery, partially due to fear of awake surgery. On the other, exact implantation in the DBS motor domain target necessitates the use of a neural navigation system that relies on awake brain signals. Centers that do not use a navigation system depend on pre-operative MRI and CT scans which can often not reflect accurate brain alignment due to image distortion, fusion errors and brain shift^34–36^. This may lead to incorrect placement of the electrode^37–39^.

Our findings suggest that an interleaved propofol-ketamine sedation protocol could be an ideal solution for three major reasons. We have shown that propofol sedation can be successfully transitioned to ketamine sedation in a timely manner, allowing practical use in the OR. Second, ketamine does not disrupt the brain’s awake activity and thus enables the use of the neural navigation system as usual. Finally, by interleaving the propofol and ketamine we can imitate the natural sleep cycle, potentially providing a more beneficial sedative state for the patient.

In general, sedatives are kept consistent across a procedure (barring unexpected events) keeping the brain state consistent as well, unlike the cyclical, staged, sleep rhythm and sedation interval routines used in intensive care units. Consequently, it is unsurprising that up to 80% of patients report waking up from sedation with a feeling of drowsiness^40^ signifying that sedation does not equate to natural sleep. Moreover, natural nocturnal sleep has been found to be markedly disturbed following surgical procedures^41^. Disturbances of sleep are highly prevalent in basal ganglia-related neurodegenerative disorders, particularly PD^42–44^. Enabling a more restful sedation could greatly benefit these patients who already suffer from sleep difficulties.

Both sedation and the sleep cycle reversibly alter consciousness. The type of sedative or sleep stage can determine what type and level of consciousness one experiences. Contrary to previous theories that anesthesia and sleep are states in which the brain is switched off, newer studies suggest that the brain’s electrophysiology is modified in accordance with consciousness level. Further, recent research reports similar electrophysiological changes, namely modification of evoked alpha and gamma power, during ketamine sedation and REM sleep compared to propofol anesthesia and NREM sleep^45^. The authors argue that ketamine sedation and REM sleep are both states of disconnected consciousness while propofol anesthesia and NREM sleep are states of unconsciousness. These results are very much in line with our findings of high and low frequency power alterations grouping ketamine with REM sleep and propofol with NREM sleep. Further, though both the propofol and ketamine sedation recording sessions were performed at what is considered moderate sedation by the American Society of Anesthesiologists (ASA), this and other reports suggest we should be considering the above definitions of dissociated and reduced consciousness instead when discussing the electrophysiology of the brain.

Though ketamine was highly touted for its cardiorespiratory preservation properties after its invention in the 1960’s, over the years it has become less commonly used due to the view that it increased salivation and upper airway secretions, its rise in popularity as a drug of abuse, and the fear of instigating frightening hallucinations^46^. Hallucinations do seem to occur in up to 50% of adult sedations under ketamine^47^, suggesting that limited use is warranted. However, the content of the hallucinations might be context dependent. Further, the power of suggestion could potentially affect the content of the hallucination and its connotations^48^. Ketamine hallucinations have also been reportedly blocked by prior or co-administration of GABA agonists, like propofol, in humans undergoing surgical procedures^49,50^.

Propofol is the most commonly used sedative in the OR today and for good reason, it is fast acting and effective. These properties make it an excellent choice for surgeries with aspects that might be anxiety-provoking. Additionally, procedure related pain control may be accomplished by systemic analgesic medication administration, and performance of local anesthesia infiltration and/or supplemented nerve blocks as done routinely nowadays. Our IPK protocol harnesses the positive properties and concerns of both propofol and ketamine to deliver the best experience for the DBS patient and surgeon/electrophysiologist. We propose propofol be used during the scalp incision and burr-hole creation due to the fearsome nature of this stage of surgery. Then, the lights in the OR should be turned down and the staff should make an effort to create a calm environment while ketamine is administered (mimicking the conditions in our dark and noise attenuated recording room). Ketamine sedation should continue for the short period of neural navigation, limiting the time in which hallucinations can occur. Once navigation has been completed, the patient should be switched back to propofol for the incision and burr-hole creation in the second hemisphere. We believe this addresses both main concerns by minimizing hallucination possibility through limited time exposure and creating a relatively relaxed context while still enabling the use of neural navigation.

While our study gives a solid starting point towards asleep DBS, it does have some limitations. First, African Green Monkeys share a similar dopaminergic system that of humans and therefore it is reasonable to use them as a model for Parkinsonism and to record from the basal ganglia. However, they are still models so no absolutely conclusive answer can be given regarding humans. Second, as commonly done and recommended in NHP studies our study had a small sample size of 2 NHPs^51^. Third, the parkinsonian NHPs in this study were only treated with dopamine replacement therapy for a short time prior to recording while in human DBS patients they are usually treated with dopamine replacement therapy for many years prior to surgery. We cannot discount that the effects of chronic dopamine therapy on electrophysiological properties and recordings. Thus, to test our proposed IPK sedation protocol in humans a comprehensive clinical study should be conducted. The current study should be carefully followed by prospective human study that hopefully would lead to paradigm change in DBS practice enabling high-quality physiological navigation during asleep DBS.

## Methods

The study’s ethical permission number is MD-18-15449-5. The Hebrew University of Jerusalem is an Association for Assessment and Accreditation of Laboratory Animal Care (AAALAC) internationally accredited institute.

### Surgery

Surgical preparation of the NHP included two procedures carried out under general and deep anesthesia (Isoflurane and N_2_O inhalation anesthesia, induction by IM 0.1 mg/kg Domitor and 10 mg/kg Ketamine) and in aseptic conditions. The first surgery consisted of a craniotomy and implantation of a chamber, head holder and EEG electrodes. The second surgery involved the implantation of a subcutaneous ported central venous catheter. Both surgeries were carried out by board-certified surgeons (ZI and VK). Anesthesia and perioperative treatment was supervised by an experienced anesthesiologist (SF) and the veterinary team of The Hebrew University of Jerusalem.

During the first surgery a 27×27 mm polyetherimide (PEI, MRI compatible, Alpha-Omega, Nof HaGalil, Israel) recording chamber and a head-holder were surgically implanted in the animal’s skull (see details in previous publications of the lab^52–54^). The chamber was positioned in a suitable location that allowed electrophysiological recordings from the frontal cortex and external segment of the globus pallidus (GPe). The location of the chamber was determined according to a primate stereotaxic atlas^55^. During the surgery, six EEG screw electrodes were positioned bilaterally, in the frontal, parietal and occipital positions of each hemisphere in the NHP’s skull.

In the second surgery, a subcutaneous ported vascular catheter (port-a-cath, Medcomp, PA, USA) was placed. Procedure for insertion of a port includes formation of a subcutaneous pocket for the port, fixation of the port to fascia by sutures, tunneling of the vascular catheter and cannulation of the superior vena-cava using Seldinger’s Technique. The procedure was carried out under ultra-sound imaging, and the final location of the catheter was verified by X-ray.

In addition to general anesthesia, the area of the surgical incisions was infiltrated with local anesthetic (2 ml of Bupivacaine 0.25%). Peri-operatively, the animal was treated with antibiotics (Ceftriaxone 35 mg/kg, PanPharma, Luitré-Dompierre, France), steroids (Dexamethazone 0.5 mg/kg, Kern Pharma, Barcelona, Spain) and analgesic (Optalgin 20 mg/kg, Teva, Petach-Tikva, Israel) medicine. It was allowed at least 4-5 days of recuperation after surgery before resuming training and experimental work.

To maintain patency of the port and intravascular tubing heparin (2.5-3 ml, 100 IU/ml, Bichsel, Interlaken, Switzerland) was used to flush the port after recording session and at least once weekly when no recordings took place. The cranial chamber was flushed every other day (including before and after recording) with a saline-neomycin solution, and dura scrapping was done several times (every 4-6 weeks) under IM Domitor (0.1 mg/kg) and Ketamine (10 mg/kg) sedation to enable penetration of the dura by microelectrodes.

### MRI Imaging

Following recovery from surgery, the NHPs underwent a 3T MRI examination (Figure 1a) to verify correct placement of the chamber and to determine its precise stereotaxic location (see details in previous publications of the lab^32^). The imaging procedure was carried out while the NHP was sedated with IM Domitor (0.1 mg/kg) and Ketamine (10 mg/kg).

### Sedation Procedure

#### Recording session protocol

The NHPs were living in an open yard with companions when there are no sedation requirements. Recording sessions were performed approximately once every 2-3 days in a pseudo-randomized order. Each session consisted of a saline baseline, sedation, and saline washout period, each lasting one hour. Sedation agents used were propofol (Raz Pharmaceutics, Kadima Zoran, Israel or B Braun, Melsungen, Germany), ketamine (Vetoquinol, Lure Cedex, France), dexmedetomidine (Kalceks, Riga, Latvia), remifentanil (Mylan, Canonsburg, PA) and scopolamine (Sterop, Brussels, Belgium). For IPK session, 40min-20min-40min-20min-20min of propofol-ketamine-propofol-ketamine-propofol (without last propofol for IPK sessions of monkey Ch before MPTP-treatment) were administered between one-hour saline and one-hour saline washout. Here, we report only on ketamine and propofol effects.

#### Fasting

The evening prior to sedation the NHP was taken from the open yard where she is normally group housed and placed, usually with one of her peers, into a clean cage with no food and ad-libitum water. Sedation sessions were carried in the day after. Two to three hours before sedation administration the water access was closed. During the sedation session vital signs (heart and respiration rate, end-tidal CO_2_) were monitored (Figure 1c). Pulse oximetry and non-invasive blood pressure measurements were found to be less reliable in our setup and were not used in this study. Preset limits for ending sedation (e.g., end-tidal CO_2_ >60 mmHg) were never reached in this study. Following the conclusion of the experimental session the monkey was again placed in a clean cage. Then, at ∼45 minute intervals, and depending on their clinical status, the water access was opened followed by a bell pepper and finally their normal pellet food. The NHP’s were given at least 2-3 days to rest and recover between sedation sessions.

#### Titration procedure

In order to ensure that the monkey was at a consistent moderate level of sedation, titration sessions were performed for each drug. In these sessions, a low infusion rate was first employed, slowly raising the rate in 15-20 minute steps until the desired level of sedation was achieved. No more than one hour of sedation was given per day. The level of sedation was based on amount of time the monkey’s eyes were closed and blink response to air puff stimulation which was directed to the eye. The air-puff was given once between 30 and 120 seconds (monkey Ch) or twice with minimal 30 second interval between 20 and 280 seconds (monkey Cs) into each 6-minutes’ time block randomly. Figure 1d shows the details of the titration process for each drug.

#### Drug Preparation

For every sedation session, sterile normal saline (NaCl 0.9%) was drawn into a 60 ml syringe and placed in the infusion pump (Injectomat Agilia, Fresenius Kabi). During the baseline period the rate of infusion was set to 3 ml/hr or 15 ml/hr (monkey Ch and Cs respectively), during sedation the total rate of the infusion (drug + saline) was 15 or 20 ml/hr (monkey Cs and Ch respectively) and during the washout the rate of saline infusion was kept at the same level as during sedation. The rate of the infusion was constant over the 1-hour periods, i.e. no target-control infusion (TCI) techniques were used.

#### Propofol

20 ml vials of 10 mg/ml (1%) propofol were drawn into a 60 ml syringe. Infusion rate during sedation sessions was set according to the NHP’s weight, i.e. 3 or 4.8 ml/hour, approximately 120 or 190 mcg/minute/kg for 4.2 kg NHP.

#### Ketamine

0.5 ml of 100 mg/ml ketamine was mixed with 49.5 ml normal saline in a 60 ml syringe. Infusion rate during sedation sessions was set according to the NHP’s weight, i.e. 10.1 ml/hour, ∼40 mcg/minute/kg for 4.2 kg NHP.

#### Port-a-cath protocol

Before every recording, the skin around the port was cleaned several times with chlorhexidine 2% in ethanol 70% solution. A sterile field was prepared. A non-coring needle (MiniLoc, Bard, Salt Lake, USA) was inserted into the proper puncture site. The dead space of the port and vascular catheter (previously locked with heparin) was drawn and the port’s patency checked by flushing it with 5 ml of normal saline. At the end of each of the recording sessions, the port was locked with 3 ml of heparin (100 IU/ml). The non-coring needle was removed and the skin covering the port was sterilized with chlorhexidine 2% in ethanol 70% solution. The hair around the port area was clipped roughly every week.

### Induction of Parkinsonian Symptoms

The NHPs were treated with MPTP-hydrochloride (Sigma, Israel) in a negative pressure isolated room to induce parkinsonism. Five IM injections of 0.35 mg/kg/injection were made over the course of four days (two injections on the first day) under ketamine (10 mg/kg IM) sedation. The NHP was moved back to their previous room 72 hours after last MPTP injection. Before, during and following induction of parkinsonism a modified Benazzouz primate parkinsonism scale^56^ was used to assess symptoms severity level (Figure 1b).

One week following the first MPTP injection, the NHPs were severely akinetic and unable to feed themselves. The experimenters fed the NHPs twice a day, seven days a week with using a pediatric nasogastric tube. A single feeding dose equals ∼30 cc water and 50-70 cc of Ensure Plus (Abbott Laboratories, Abbott Park, Illinois), a high calorie (1.5 Kcal/cc) nutritional shake. The dopamine replacement therapy 1/8∼1/2 tablet, 250 mg L-DOPA and 25 mg carbidopa per tablet (Dopicar, Teva Pharmaceutical Industries, Israel) was administered with every nasogastric feeding. Neuronal recordings during the parkinsonian condition began when the animal exhibited severe parkinsonian symptoms approximately 7 days after the first injection. Twelve hours prior to recording sessions, dopamine replacement therapy was stopped to allow for washout. The Parkinsonian symptoms were stable all over the recording period (Figure 1b, top subplot).

### Experimental Set-up

The experiments were performed using two separate rooms. 1. An experimental room, where vital signs (Nasal End-tidal CO2, respiratory rate, ECG and heart rate) were monitored (Mindray, BeneVision N12, Shenzhen, China) and the experimenters manipulated the infusion pumps, the electrode vertical positions, operated the data acquisition tools and performed on-line analysis (e.g. spike sorting). 2. A noise attenuated (Industrial Acoustics Company, Controlled Acoustic Environment, IL, USA) recording room, in which the NHP was located. The recording room was dark during the recording and infra-red cameras (Metaphase Technologies Inc., PA, US) were used to monitor the NHP. Live video (Figure 1a, d-g, ImagingSource, NC, USA) recording was collected at 50 Hz during all sessions allowing the researchers to monitor the animal and determine whether the NHP’s eyes were open or closed. This was then synchronized with the neural data collected using AlphaLab SnR (Alpha-Omega Engineering, Nof Hagalil, Israel). Figure 1a depicts a schematic diagram of the experimental setup.

### Polysomnography

We recorded the electrooculography (EOG) signal bilaterally using disposable subdermal needle electrodes (Rhythmlink, Columbia, US). We placed one electrode 1 cm below the left outer canthus, and another electrode 1 cm above the right outer canthus. We recorded the electromyography (EMG) signal from the NHP’s trapezius muscle contra-lateral to the port by two disposable subdermal needles placed ∼1cm apart. We continuously recorded the surface electroencephalogram (EEG) signal, using the chronic EEG electrodes that were implanted during the surgery. Six electrodes were used – two frontal, two parietal and two occipital. Cranial implant fastener (slip under the skull and held in place with a double nuts, 6-YCI-06L, Crist Instruments, Maryland, USA) was used as a ground and a reference. The EOG, EMG and EEG were sampled at 2750 Hz. Detailed polysomnography methods can be found here^44^.

### Behavioral Measures

To model the physical touch that is often used to assess a patient’s sedation level (eyelash reflex), we measured the blink evoked by a short duration air-puff directed at the eyes of the NHP using computer-controlled solenoid valve (SD Instruments, CA, USA) connected to a pressurized gas tank. The air-puff was administered randomly in time with a pressure of 8 bars and duration of 0.2 seconds.

### Electrophysiological Recording Procedures

During recording sessions, the NHP’s head was immobilized by a head-holder. Then, eight independently controlled glass-coated tungsten microelectrodes (impedance 0.3-1.2 MΩ at 1000 Hz) were advanced into the brain (EPS; Alpha-Omega Engineering, Nof Hagalil, Israel. Smallest step, 1 µm) towards the target regions^57^. Raw signal (0.1-9000 Hz), spiking activity (SPK, 300-9,000 Hz) and local field potential (LFP, 0.1-300 Hz) were recorded. Cells were selected for recording as a function of their isolation quality and optimal signal-to-noise ratio. Online, the researchers monitored the quality of the cells and noted defining characteristics, discharge rate and a letter-grade rating of overall quality of the recorded spikes. The data was continuously sampled at a frequency of 44 KHz (raw and SPK data) and 1375 Hz (LFP data) by 16 bits analog to digital converter (SnR, Alpha-Omega). Offline, we used the isolation score as a criterion to define an appropriate unit database for subsequent analyses^58^.

## Data Analysis Methods

All data analysis was conducted using in-house Matlab (MathWorks, Natick, MA; version R2018b) scripts.

### Eye Closure Analysis

For each sedation session and sleep night session, a modified version of eye open/close detection tool^59^ was used. An area of interested (AOI) was selected to cover the pixels of two eyes and applied to frames of the whole session (nights). The averaged grayscale and 1st principal component of AOIs were used to form a 2-D space, and 2-D threshold were manually selected to define eye open/close of frames.

### Air Puff Stimulation Analysis

For each sedation session, the video frames of saline (all frames), sedation (5 min after sedation initiation to washout) and washout (5 min after washout initiation to end) were used to calculate eye closure proportion for each stage. Frames were aligned to the end of air-puff to show eye closure rate around air-puff for each stage. Sequential eye closed frames which were longer than 1 second were marked as long duration closure frames, otherwise they were marked as short duration closure (blink) frames. The same eye closure proportion analysis was applied to all, short and long duration eye-closures.

### Sleep Staging

To determine the sleep stages, the polysomnography measurements of EEG (filtered 0.1-35 Hz), EOG (filtered 0.1-35 Hz), EMG (filtered 10-100 Hz) and eye state (open/closed) were used. Sleep staging was done using a semiautomatic staging algorithm (custom software) which took 10 second non-overlap epochs and clustered them based on three features: (i) the average ratio of high/low EEG power across all contacts. The average power at 15-25 Hz (related to waking) was divided by the average power at 0.1-7 Hz (related to sleep); (ii) root mean square (RMS) of the EMG signal; and (iii) eye-open fraction (open/all). Every 10 second epoch was represented as a point in three-dimensional feature space, usually forming three clusters: for wakefulness (high EMG RMS, increased EEG high/low ratio, eye-open fraction close to 1), NREM (low EMG RMS, decreased EEG high/low ratio, eye-open fraction close to 0), and REM (very low EMG RMS, increased EEG high/low ratio, eye-open fraction close to 0). Before semiautomatic clustering, a trained expert manually clustered 10∼30% of the night epochs based on EEG, EMG, EOG and eye open/close. The staging results provided by the semiautomatic algorithm were accepted for further analysis only if they matched the expert staging in more than 85% of the tested epochs. Two nights (monkey Ch) which EMG RMS has no clear separation between REM and NREM were removed from the database. For further elaboration on sleep staging analysis see here^60^.

### Spectrogram Analysis

For each signal type (e.g. EEG, local field potential (LFP), spiking activity (SPK), etc.), Welch’s power spectral densities as a function of time (spectrogram) with 12 second window, 6 second overlap, frequency range 0.5 Hz to 100 Hz with 0.5 Hz resolution was calculated by 60 second moving window with 30 second step. DC (direct current, 0 Hz) was removed for each 60 second window by subtraction of the window mean. The SPKs were rectified to capture the low-frequency oscillations of the discharge rate^61^. To reduce the noise at specific frequencies, power densities of 48-52 Hz (power line frequency), 60 Hz, 99-100 Hz were linearly filled with power densities of flanking frequencies. Power densities of 16.5-18 Hz were linearly filled for Ctx/GPe LFP. The power densities for certain time bins were set to not-a-number (NaN), if the electrode position was not stable (for LFP/SPK) or total power densities were defined as outliers by more than 1.5 interquartile ranges above the upper quartile or below the lower quartile for consequential 15 bins (8 min). The power densities of each time bin in the spectrogram were normalized to 1. Then the proportion of power density for each frequency were normalized to z-score of saline periods (Figure 1h, top subplot). The normalized spectrograms were averaged within the same signal types.

Similar analysis was applied for sleep periods longer than 60 seconds. The power densities were set to NaN, if the electrode position is not stable (for LFP/SPK) or total power densities were defined as outliers of segments of the same sleep stages. After normalization of total power in each time bin to 1, the proportion of power density for each frequency were normalized to z-score of wake periods. The normalized spectrograms of wake, NREM and REM were aligned to its own initiation and averaged within same signal types.

### Coherogram Analysis

For each pair of signal types (e.g. EEG-EEG, Ctx LFP-GPe SPK, etc.), magnitude-squared coherence (MSC) as a function of time (coherogram) with 12 second window, 6 second overlap, frequency range 0.5 Hz to 100 Hz with 0.5 Hz resolution were calculated by 60 second moving window with 30 second moving step. DC (direct current, 0 Hz) was removed for each pair for each 60 second window by subtraction of the window mean of each signal. The SPK signal was rectified to capture the low-frequency oscillation of discharge rate, and sampling rate of pairs of signals was down sampled to the sampling rate of the signal with lower sampling rate. To reduce the noise at specific frequencies, LFP related MSC of 16.5-18 Hz, 48-52 Hz, 60 Hz, 99-100 Hz were linearly filled with MSC of the flanking frequencies. EEG related MSC of 48-52 Hz, 60 Hz, 99-100 Hz were similarly linearly filled. The MSC for certain time bins were set to NaN, if the electrode position was not stable (for LFP/SPK related MSC) or total MSC was defined as an outlier by more than 1.5 interquartile ranges above the upper quartile or below the lower quartile for more than 15 bins (8 min). MSC values were bounded between zero and one and therefore are not linearly distributed. We used the Fisher z-transform to normalize the MSC distribution. The Fisher z-transferred squared root of MSC of each time bin in coherogram were normalized to 1. Then the proportion for each frequency was normalized to z-score of saline periods (Figure 1h, lower subplot). The normalized coherograms were averaged within same pairs of signal types.

Similar analysis was applied for sleep periods which were longer than 60 seconds. The MSC was set to NaN, if the electrode position was not stable (for LFP/SPK related MSC) or total MSC was defined as outlier of segments of the same sleep stages. The MSC were normalized by frequency and normalized to z-score of wake periods. The normalized coherograms of wake, NREM and REM were aligned to its own initiation and averaged within the same pair of signal types.

### High/low power/synchronization difference

For each spectrogram, the normalized power of 12-40 Hz was averaged to represent the power of the high frequency domain and the normalized power of 0.4-4 Hz was averaged to represent the power of the low frequency domain (frequency ranges are shown by vertical lines in Figure 1h right). The difference between the normalized high frequency power and the normalized low frequency power were used to represent high/low difference of the spectrogram. For each sedation session (nights), all normalized high/low power difference were averaged within different signal types, then averaged though different signal types to represent the high/low power difference of this sedation session (nights). The same frequency ranges were used for the coherogram to calculate the high/low synchronization difference.

#### Rehabilitation

Following completion of the study, the NHP’s continued to be fed through naso-gastric tube and were treated 1-3 times daily with dopamine replacement therapy. After partial recovery of the Parkinsonian symptoms (2-3 months after the MPTP injections) the head apparatus and the port-a-cath were removed under general anesthesia, perioperative antibiotics, and analgesic treatment. As they improved, the NHPs were gradually reintroduced to their group starting with 10 minute one on one time with another NHP, then one on one overnights and during feeding times, and finally full reintroduction. After parkinsonian symptoms were deemed minor (Figure 1b, lower subplot) and the NHP showed the ability to eat and interact fairly normally within the group, they will be sent to the Israel Primate Sanctuary (www.ipsf.org.il). One has already been successfully rehoused at the sanctuary, another is still undergoing the recovery process.

## Acknowledgments

We thank Tamar Ravins-Yaish for assistance with animal care; Atira Bick and ELSC neuroimaging unit (ENU) for assistance with the MRI scan; and Sharon Freeman for general assistance. We thank Anatoly Shapochnikov for help in preparing the experimental setup. This work was supported by grants from the Israel Science Foundation, German Collaborative Research Center TRR295 (Returning dynamic motor network disorders using neuromodulation) Adelis and the Rosetrees Trust (to HBe) and research grant of the department of anesthesiology, Hadassah Medical Center (to SF).

## Author Contributions

OBY, SF and HBe conceived of the research questions and built experimental setup. JG, HBa, OBY, SF and UWR collected the data. JG, HBa and OBY performed the data analysis. HBa and HBe wrote the manuscript and JG made the figures. VK and ZI performed the necessary surgeries. HBe oversaw all aspects of the work. All authors have read, discussed and approved the final version of the manuscript.

