## Supplementary Tables and Figures for "Toward asleep DBS: Cortico basal-ganglia neural activity during interleaved propofol/ketamine sedation mimics NREM/REM sleep activity"

| Normal | | | | | | | | | | | | | | |
| --- | --- | --- | --- | --- | --- | --- | --- | --- | --- | --- | --- | --- | --- | --- |
|  | Propofol | | | Ketamine | | | IPK | | | Sleep | | | | |
|  | Recording Sessions | Ctx sites | GPe sites | Recording Sessions | Ctx sites | GPe sites | Recording Sessions | Ctx sites | GPe sites | Recording Nights | | Ctx sites | | GPe sites |
| Monkey Ch | 9 | 22 | 20 | 8 | 26 | 18 | 1 | 4 | 3 | 6 | | 18 | | 12 |
| Monkey Cs | 8 | 18 | 20 | 6 | 17 | 14 | 2 | 5 | 7 | 13 | | 34 | | 33 |
| Total | 17 | 40 | 40 | 14 | 43 | 32 | 3 | 9 | 10 | 19 | | 52 | | 45 |
| MPTP treated | | | | | | | | | | | | | | |
|  | Propofol | | | Ketamine | | | IPK | | |  | | | | |
|  | Recording Sessions | Ctx sites | GPe sites | Recording Sessions | Ctx sites | GPe sites | Recording Sessions | Ctx sites | GPe sites |  |  | |  | |
| Monkey Ch | 3 | 11 | 11 | 3 | 12 | 8 | 2 | 7 | 6 |  |  | |  | |
| Monkey Cs | 3 | 7 | 8 | 3 | 8 | 10 | 2 | 6 | 8 |  |  | |  | |
| Total | 6 | 18 | 19 | 6 | 20 | 18 | 4 | 13 | 14 |  |  | |  | |

**Table.S1 | Recording database.**

For each sedative drug and sleep, recording sessions, nights and sites from both monkeys are given. Abbreviations as in Fig.1.

|  | Propofol | | | | | |
| --- | --- | --- | --- | --- | --- | --- |
|  | Normal (N=17) | | | MPTP (N=6) | | |
|  | Sal-Ppf | Ppf-Wash | Sal-Wash | Sal-Ppf | Ppf-Wash | Sal-Wash |
| All | *** 0.0004 | *** 0.0004 | 0.36 | * 0.0312 | * 0.0312 | 0.44 |
| Long | *** 0.0004 | *** 0.0004 | 0.33 | * 0.0312 | * 0.0312 | 0.44 |
| Short | *** 0.0003 | *** 0.0003 | 0.62 | 0.0625 | * 0.0312 | 0.31 |
|  | Ketamine | | | | | |
|  | Normal (N=14) | | | MPTP (N=6) | | |
|  | Sal-Ktm | Ktm-Wash | Sal-Wash | Sal-Ktm | Ktm-Wash | Sal-Wash |
| All | *** 0.0001 | *** 0.0001 | ** 0.0040 | * 0.0312 | * 0.0312 | * 0.0312 |
| Long | *** 0.0001 | *** 0.0001 | ** 0.0067 | * 0.0312 | * 0.0312 | * 0.0312 |
| Short | 0.95 | ** 0.0012 | * 0.0245 | 0.44 | 0.56 | 0.22 |

**Table.S2 | Eye closure proportion changed during propofol and ketamine sedation.**

Statistics of Fig.1e raster and bar plot. Two propofol sedations before MPTP with no air puff applied were excluded from rate plot in Fig.1e. P-value is given, Wilcoxon signed-rank test. Abbreviations as in Fig.1.

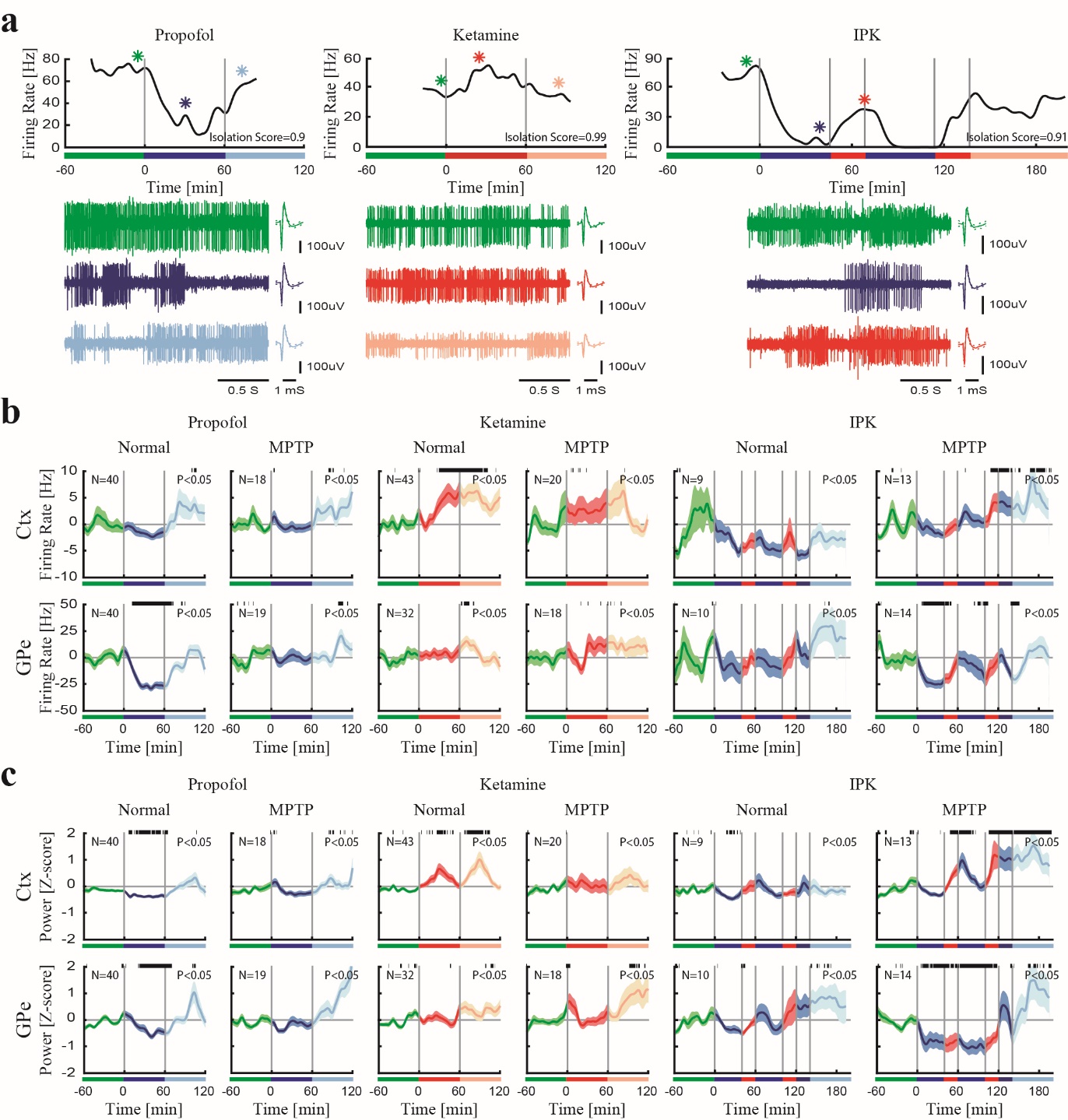

**Fig.S1: Firing properties change during ketamine and propofol in cortex and the basal ganglia. a.** Examples of firing rate of GPe during propofol (left), ketamine (center) and IPK (right) sedation sessions. The spiking from time locations marked with asterisks is shown (lower). **b.** Averaged firing rate difference of multi-units of Ctx (upper) and GPe (lower) during propofol (left), ketamine (center) and IPK (right) before and after MPTP-treatment. Top black bar shows significant difference in firing rate compared to saline (p<0.05, Wilcoxon rank sum test). **c.** Normalized total power (0.5-100 Hz) of SPK of Ctx (upper) and GPe (lower) during propofol (left), ketamine (center) and IPK (right) before and after MPTP-treatment. Top black bar shows significant difference compared to saline (p<0.05, Wilcoxon rank sum test). Color represents session epochs: saline baseline (green), propofol sedation (blue), ketamine sedation (red) and saline washout (propofol, cyan; ketamine orange). Abbreviations as in Fig.1.

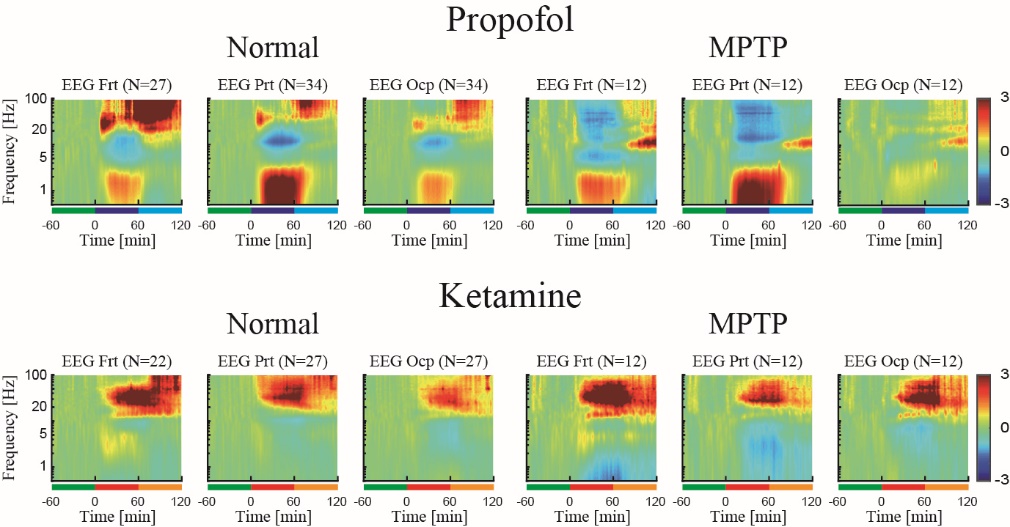

**Fig.S2: Propofol and ketamine increase low and high frequency power, respectively, in frontal, parietal and occipital EEG.** The normalized power spectrograms of Frt, Prt and Ocp EEG during propofol (upper) and ketamine (lower) before (left) and after MPTP-treatment (right). Lower bar represents 1-hour time periods of saline baseline (green), propofol sedation (blue), ketamine sedation (red) and saline washout (propofol, cyan; ketamine, orange). Number of sites is given for both monkeys in each subplot. Abbreviations as in Fig.1.

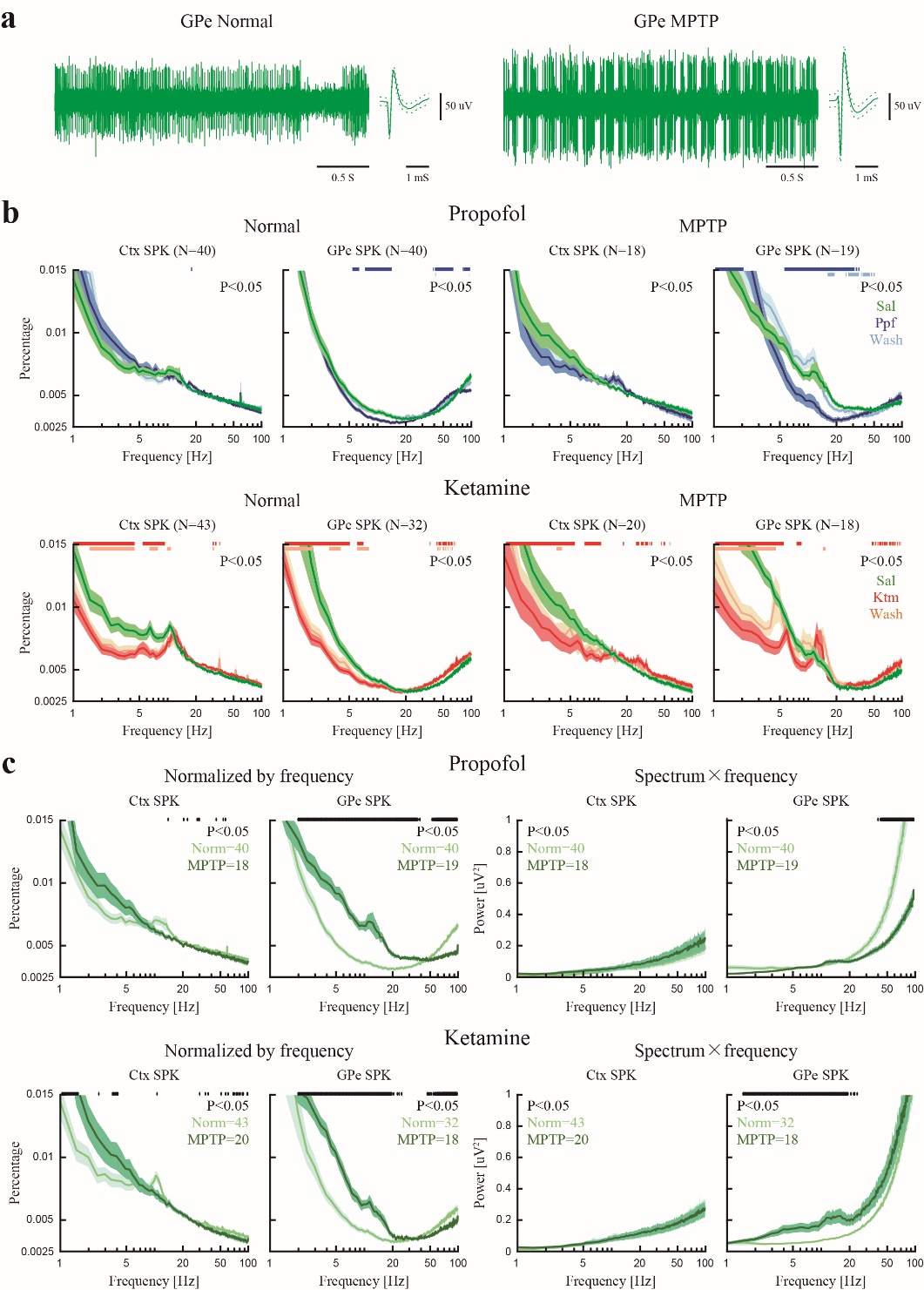

**Fig.S3: Spiking activity in the basal ganglia shows increased power in beta band after MPTP-treatment. a.** Examples of GPe SPK before (left) and after (right) MPTP-treatment. **b.** Average power spectrum densities of Ctx/GPe SPK during propofol (upper) and ketamine (lower) before (left) and after (right) MPTP-treatment. First 15min of each sedation stage is not included in averaging. Power is given as fraction of total power in the range of 0.5-100 Hz. Top color bar shows significant difference between sedation, washout and saline (p<0.05, Wilcoxon rank sum test). **c.** Averaged power spectrum densities of Ctx/GPe SPK during saline before and after MPTP-treatment. Left, normalized by total power. Right, normalized by frequency multiplication (whitening, 1/f normalization). Top black bar shows significant difference (p<0.05, Wilcoxon rank sum test). Color represents before (light green) and after (dark green) MPTP-treatment. Number of sites is given. Color codes are the same as Fig.S1. Abbreviations as in Fig.1.

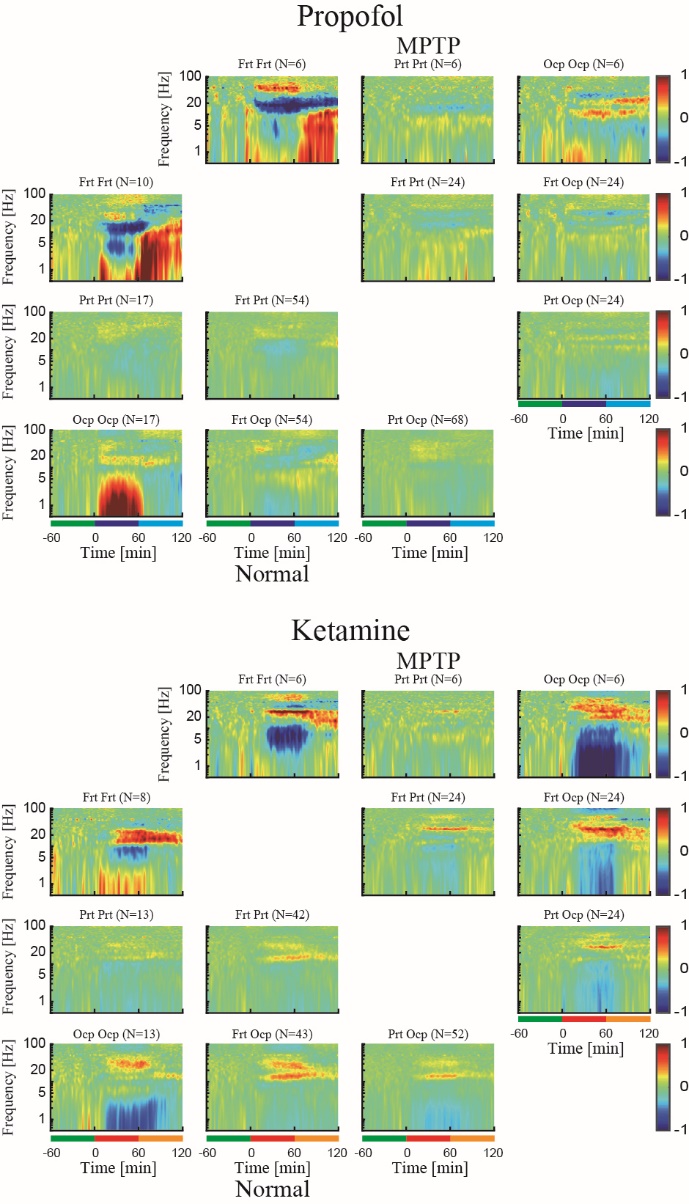
**Fig.S4: Propofol and ketamine increase and decrease low frequency EEG pairwise synchronization, respectively.** The normalized magnitude-square coherograms of all pairs of EEG during propofol (upper) and ketamine (lower) before (left) and after (right) MPTP-treatment. Lower bar represents 1-hour time periods of saline baseline (green), propofol sedation (blue), ketamine sedation (red) and saline washout (propofol, cyan; ketamine, orange). Number of pairs is given in each subplot. Color scale represents z-score. Abbreviations as in Fig.1.

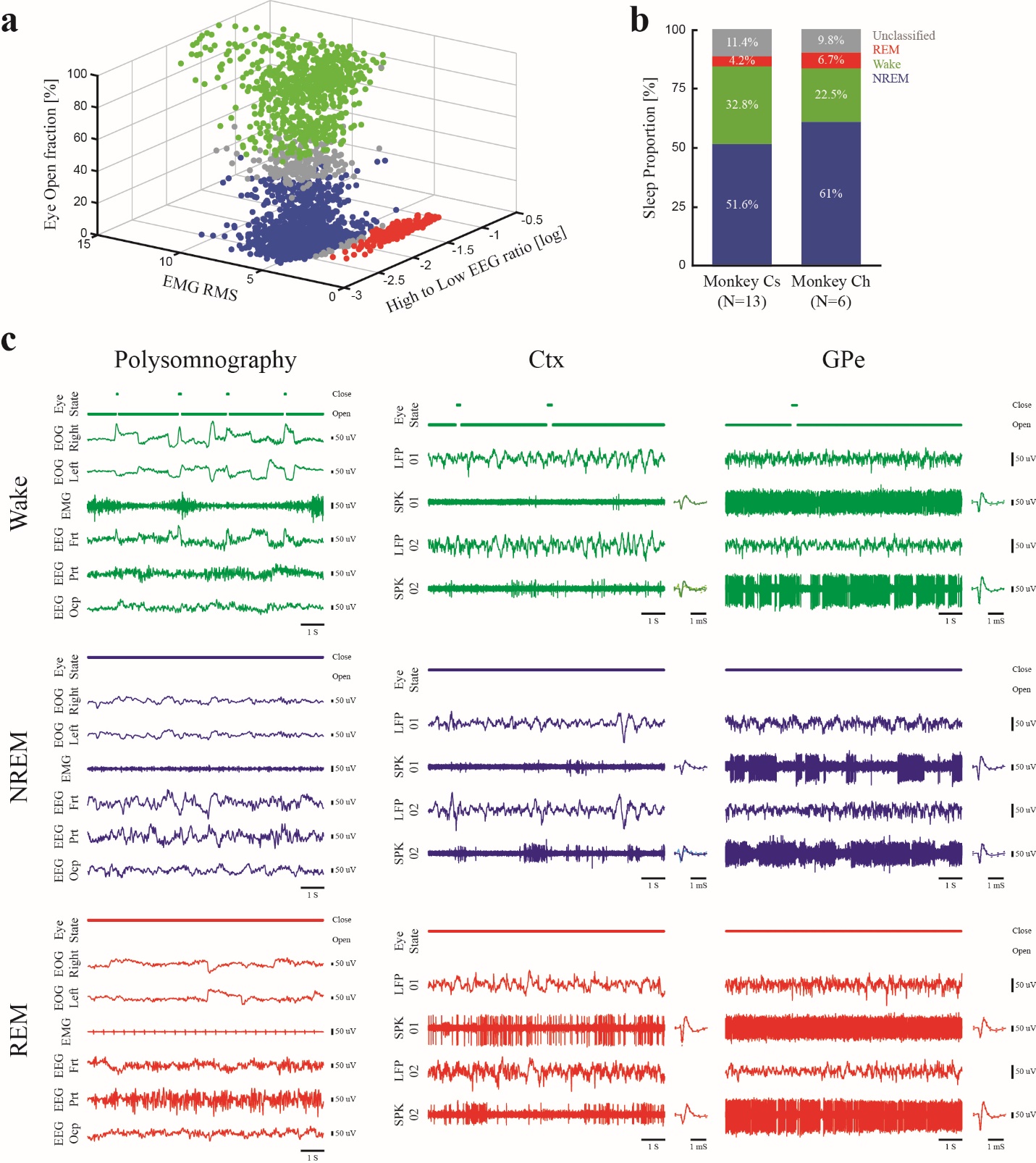

**Fig.S5: Example of sleep stage and sleep statistics. a.** One-night example of the output of the semiautomatic sleep staging algorithm (monkey Cs). The different sleep stages (wake, NREM, and REM) are staged by high/low EEG power ratio, EMG RMS, and eye-open fraction. **b.** The average proportion of sleep stages out of the all nights’ duration for the two monkeys. N is number of nights. **c.** Examples of polysomnography (left), LFP/SPK of Ctx (center) and GPe (right) during wake (upper, green), NREM (center, blue) and REM (lower, red) sleep. Abbreviations as in Fig.4.

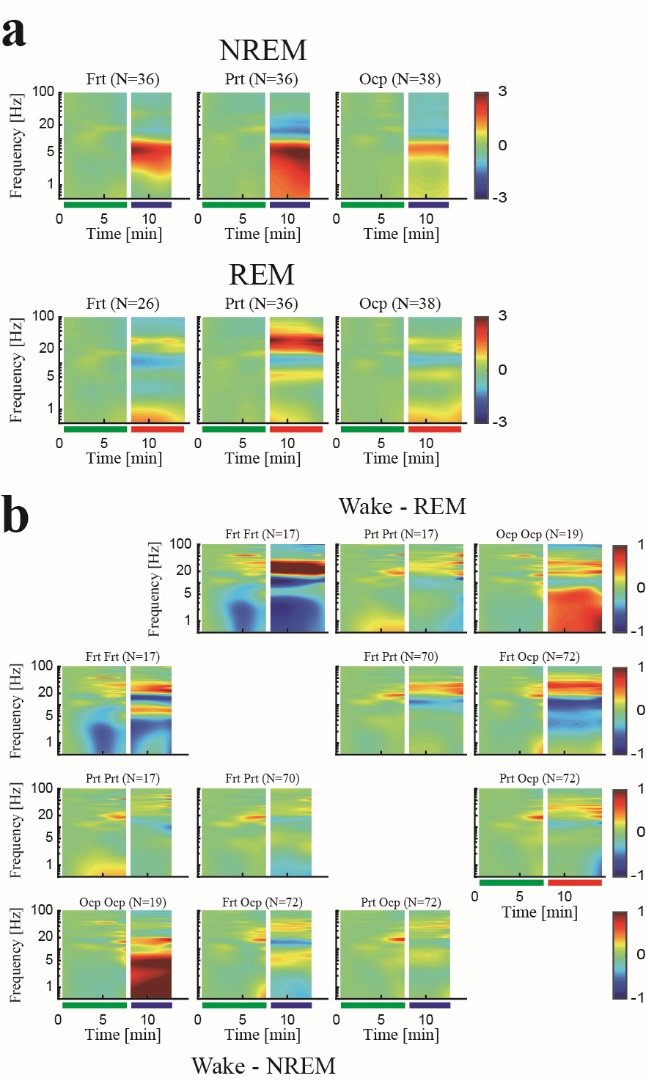

**Fig.S6: Frontal, parietal and occipital EEG during natural NREM and REM sleep show increased low frequency power/synchronization and increased high frequency power/synchronization, respectively. a.** The normalized power spectrograms of Frt, Prt and Ocp EEG during NREM (upper) and REM (lower) sleep. Lower bar represents time periods of wake (green), NREM (blue) and REM (red). **b.** The normalized coherograms of all pairs of EEG during NREM (lower left) and REM (upper right). Lower bar represents time periods of wake (green), NREM (blue) and REM (red). Number of sites (A) and pairs (B) is given for both monkeys for each subplot. Abbreviations as in Fig.4.

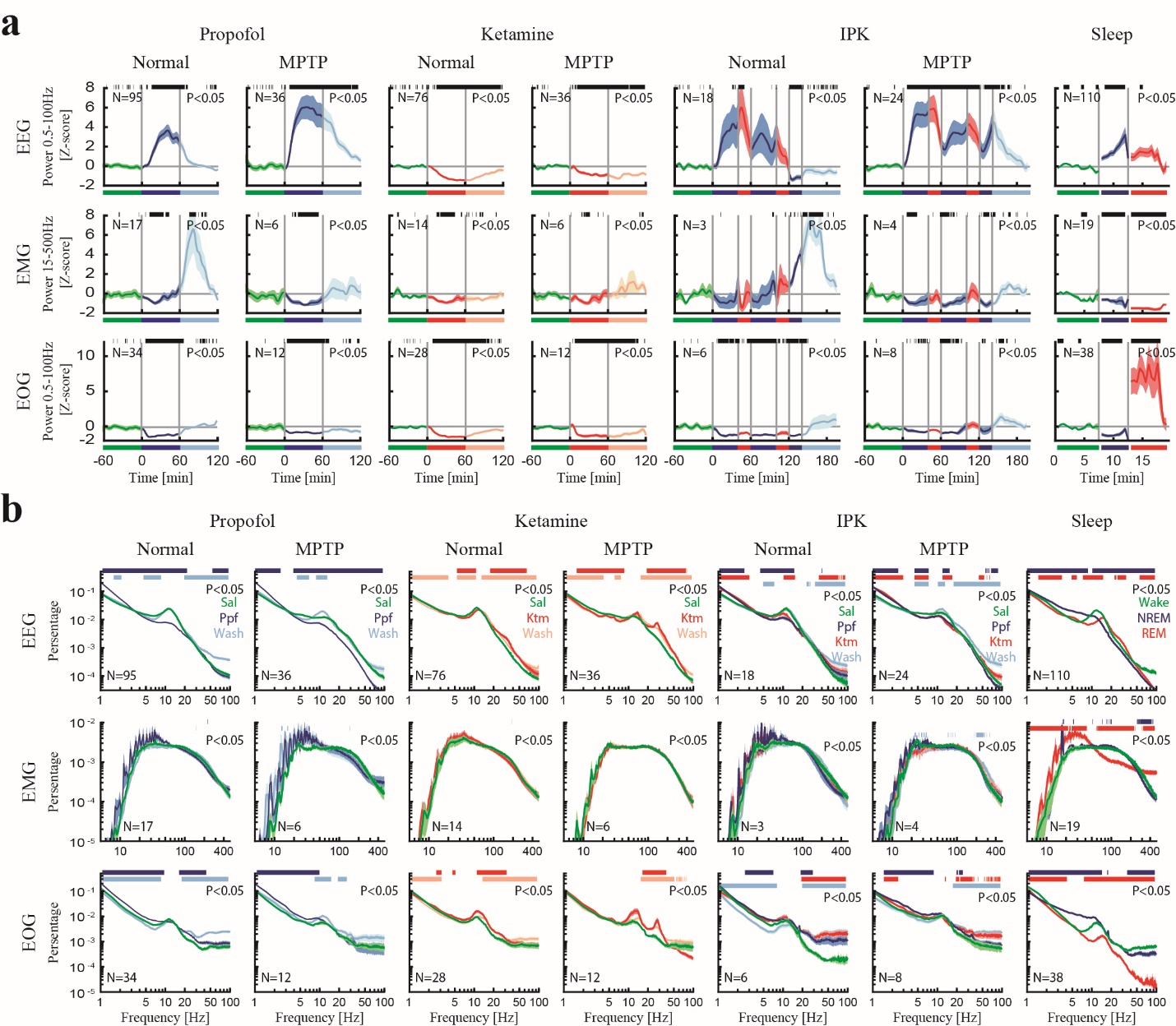

**Fig.S7: Polysomnography behaves differently during ketamine, propofol sedation and sleep stages. a.** Normalized total power of EEG (upper), EMG (center) and EOG (lower) during propofol, ketamine, IPK sedation and sleep. Top black bar shows significant difference compared to saline (wake) (p<0.05, Wilcoxon rank sum test). **b.** Average power spectrum densities of EEG (upper), EMG (center) and EOG (lower) during propofol, ketamine, IPK sedation and sleep. First 15 min (10 min for IPK) of each sedation stage is not included in the average. Power is given as fraction of total power. Top color bar shows significant difference between sedation, washout and saline or NREM, REM and wake (p<0.05, Wilcoxon rank sum test). EMG is filtered 15-1000 Hz. Color codes and abbreviations are as Fig.1 and Fig.4.

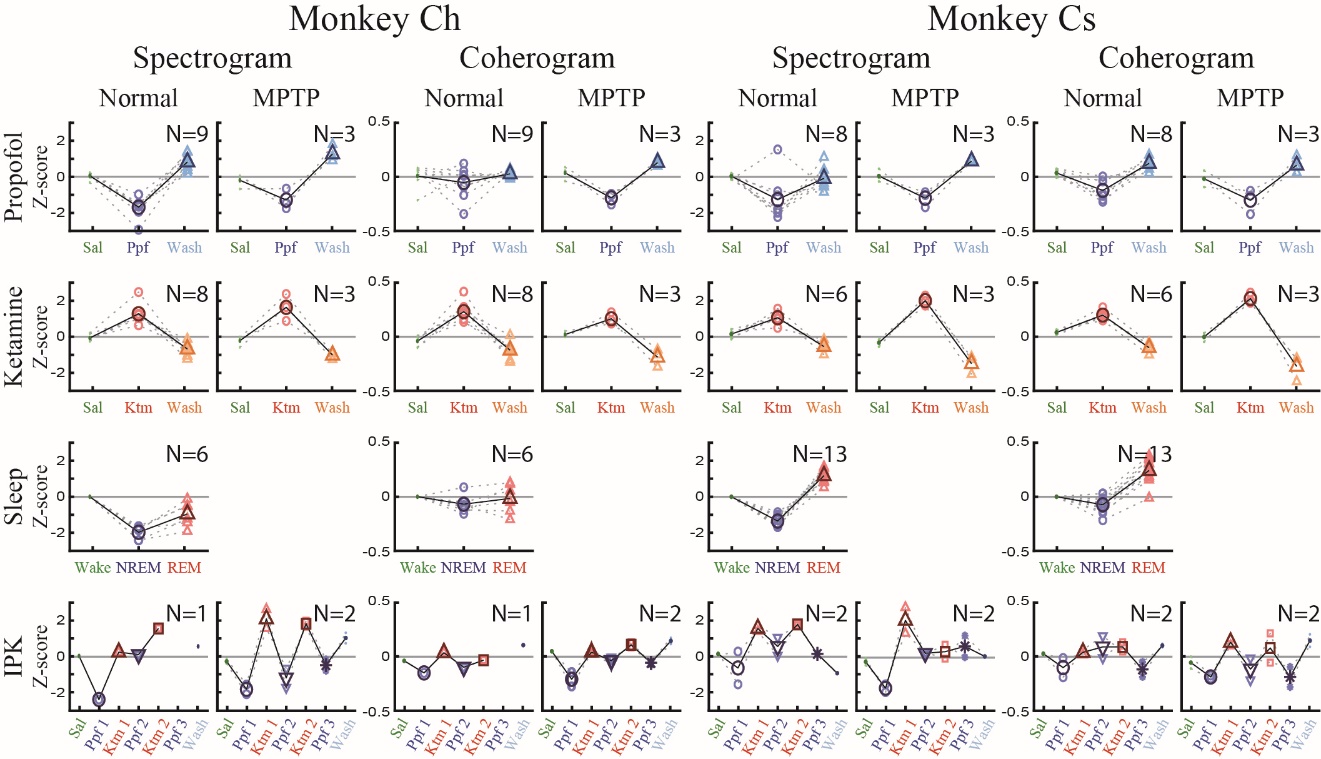

**Fig.S8: Propofol/NREM and ketamine/REM show decreased and increased high/low power/synchronization difference, respectively, in both monkeys.** Color represents saline baseline (green), propofol sedation (blue), ketamine sedation (red), saline washout (propofol, cyan; ketamine, orange), wake (green), NREM (blue) and REM (red). Number of sedation sessions or nights is given for each monkey for each subplot.
